# Mutational bias shapes protein evolution across RNA viruses

**DOI:** 10.64898/2026.07.07.737047

**Authors:** Bogdan Efimenko, Alexandr Voronka, Victoriya Skripskaya, Valeria Timonina, Alexey Agranovsky, Valerian Yurov, Konstantin Khrapko, Jacques Fellay, Konstantin Gunbin, Konstantin Popadin

**Affiliations:** Center for Mitochondrial Functional Genomics, Immanuel Kant Baltic Federal University, Kaliningrad, Russian Federation; Limnological Institute, Siberian Branch of the Russian Academy of Sciences, Irkutsk, Russian Federation; School of Life Sciences, Ecole Polytechnique Fédérale de Lausanne, Lausanne, Switzerland; Swiss Institute of Bioinformatics, Lausanne, Switzerland; Faculty of Biology, Lomonosov Moscow State University, Moscow, Russian Federation; Northeastern University, Massachusetts, USA; CellKinetica, Switzerland

**Author notes:** equal contribution (Bogdan Efimenko, Alexandr Voronka). equal contribution (Konstantin Gunbin, Konstantin Popadin).

## Abstract

Mutational biases can influence genome composition, but their contribution to protein evolution remains difficult to quantify. Here we utilize a nearly neutral framework that translates nucleotide mutational spectra into expected amino acid substitution patterns and equilibrium amino acid compositions. Using SARS-CoV-2 as a model system, we show that the viral mutational spectrum explains more than 50% of the variation in observed single-nucleotide amino acid substitutions and predicts the overall direction of proteome-wide amino acid composition change during the COVID-19 pandemic. The predictive power of the model varies with selection regime: effectively neutral and weakly deleterious substitutions conform most closely to the mutational expectation, whereas strongly constrained sites and mutational hotspots show larger deviations. This indicates that departures from the nearly neutral baseline provide a quantitative proxy for purifying and positive selection. Extending the analysis across 34 RNA virus species, we find that positive-sense, negative-sense and double-stranded RNA viruses differ systematically in their mutational spectra, and that these differences are associated with predictable shifts in proteome composition. The same relationship is detectable in RNA-dependent RNA polymerase sequences from more than 77,000 viral species. These results indicate that taxon-specific mutational bias contributes persistently to protein evolution across evolutionary scales.

## INTRODUCTION

Forecasting the paths of protein evolution represents a primary objective within the field of evolutionary genomics. Traditionally, this is viewed through the lens of natural selection acting upon a fitness landscape. However, the extent to which intrinsic mutational biases fundamentally channel these evolutionary paths remains a major unresolved question. The observation of universal biases in amino acid composition is well-established ^1^, traditionally attributed to the interplay between genetic code evolution, selection pressures and mutational forces. Considerable evidence suggests that nucleotide mutational spectra—the intrinsic biases in substitution types and rates—serve as fundamental determinants of amino acid landscapes across diverse species and genetic loci over tiny and vast evolutionary timescales ^2–6^. While earlier investigations provided primarily qualitative insights, contemporary methodological advances now permit a rigorous, quantitative exploration of how these spectra steer neutral and adaptive amino acid evolutionary dynamics^7–9^.

Because deep quantifying the effect of mutation on the protein sequence’s evolution requires vast sequence diversity and high mutation rates, RNA viruses serve as an unparalleled model system to observe these fundamental genomic mechanics in real-time. The unprecedented scale of genomic surveillance and phylogenetic reconstruction during the COVID-19 pandemic has enabled the characterization of high-resolution mutational spectra for various SARS-CoV-2 clades. These profiles are dominated by C>U transitions and G>U transversions, likely arising from cytosine deamination and oxidative damage associated with host-induced cellular stress ^10–13^. This mutational bias strongly influences genomic nucleotide content, favoring a higher frequency of Uracil at fourfold sites ^14–17^, and proteome amino acid content promoting residues frequently encoded by U-rich triplets ^10,18–20^.

In addition to SARS-CoV-2, the worldwide proliferation of sequence data has revealed the evolutionary dynamics of other viral species. Mutational spectra vary markedly within lineages and large taxonomic groups, including positive-sense (+ssRNA) and negative-sense (–ssRNA) RNA viruses ^13^. These changes are predominantly influenced by intrinsic traits, such as the fidelity of viral replication machinery, rather than by the host environment. This divergence in mutational spectra significantly influences nucleotide and codon composition ^17,21^, which, coupled with selection pressure, results in unique amino acid compositional biases among viral groups, suggesting that different lineages adhere to predictable evolutionary paths affected by their specific mutational biases.

Here, we systematically analyze, using a nearly neutral framework, how mutational biases guide viral protein evolution. By applying a nearly neutral model of amino acid substitutions to previously published SARS-CoV-2 mutational data^15,22^, we demonstrate that the mutational spectrum based on synonymous mutations at fourfold sites effectively captures observed amino acid substitution frequencies and changes in amino acid content throughout the pandemic. Extending this approach to diverse viral lineages using reconstructed spectra from 34 species, we show that group-specific mutational biases along with selection shape divergent macroevolutionary trajectories, serving as a principal determinant of global protein diversification across the virosphere.

## RESULTS

### 1. Modeling amino acid substitutions through a nearly neutral mutational framework

Utilizing the mutational spectrum of SARS-CoV-2 (Fig. 1A)—deemed nearly neutral as per references ^22^ and Supplementary Figure 1—predominantly featuring C>U transitions and G>U transversions, we sought to model the influence of this bias on amino acid composition under a largely neutral framework (assuming negligible selection, if any, for prevalent mutations). Considering the structure of the standard genetic code, we qualitatively predicted the major amino acid substitution paths driven by the two predominant single-nucleotide changes (Fig. 1B, hereafter referred to as the “butterfly plot”). For instance, amino acids whose codons are predominantly C/G-rich, such as Arg, Ala, Pro, Gly and Thr, are predicted to decrease in prevalence (“losers”) due to multiple codon substitutions that promote their loss. Conversely, amino acids frequently encoded by U-rich codons, such as Leu, Phe, Val, Ile, Ser, Cys and Tyr, are expected to increase (“gainers”). The fate of other amino acids is more challenging to predict qualitatively, as they are associated with rare nucleotide substitutions, or low number of codons, thus necessitating a numerical solution.

**Figure 1.**
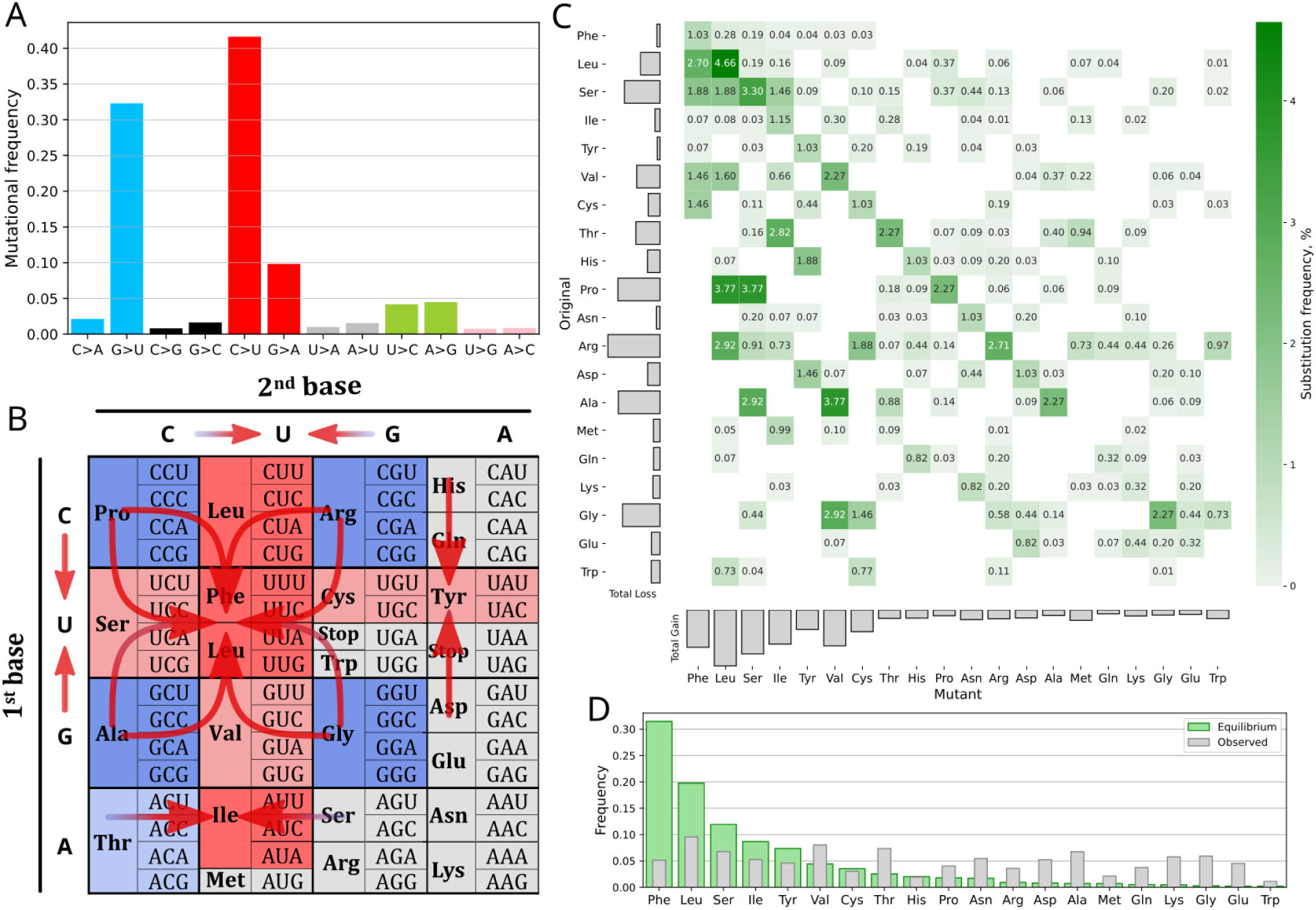
From nucleotide to amino acid spectrum of substitutions in SARS-CoV-2 under a nearly neutral model. (A) Mutational spectrum of SARS-CoV-2 (clade 20A from ^22^). (B) ‘Butterfly plot’ – reordered standard codon table with main mutational flows indicated. Red cells indicate gainers, blue cells – losers. Gray cells also indicate losers according to D. (C) Expected amino acid substitution frequencies, as predicted by the model based on the SARS-CoV-2 nucleotide mutational spectrum (clade 20A) and assuming initially uniform codon usage. Gray bars indicate the total loss (left) and gain (below) for each amino acid. (D) Comparison between the amino acid frequencies observed across the complete proteome and those predicted by the mutational equilibrium model.

To quantitatively ascertain the direction and relative pace of protein sequence evolution, we computed the anticipated spectrum of amino acid substitutions, analogous to methodologies employed in previous studies ^3,4,23,24^. For each codon, assuming equal starting frequencies, we calculated the rate of its transformation into another codon based on the virus’s specific nucleotide mutation spectrum. By summing these probabilities across codons, we predicted how often each amino acid would be replaced by another, resulting in a matrix that summarizes the expected substitution rates for all 150 possible amino acid pairs arising from single-nucleotide changes (Fig. 1C). This approach translates the mutation spectrum from the nucleotide to the amino acid level, providing a nearly neutral model for amino acid substitutions in SARS-CoV-2. Based on the spectrum of amino acid substitutions (Fig. 1C), we calculated the expected equilibrium frequencies of amino acids (Fig. 1D), which represent the theoretical attractor point in amino acid composition space. The predicted equilibrium frequencies (Fig. 1D) mirror and further clarify the butterfly plot: for instance, the seven amino acids with the highest expected frequencies (gainers) are also observable in the butterfly plot (red cells). Comparing the predicted mutational equilibrium frequencies with the observed amino acid frequencies reveals an imbalance, indicating the likely direction of future changes in amino acid composition—showing which residues would increase or decrease in frequency under conditions of weak selection pressure. Collectively, the framework described here provides a nearly neutral scenario for amino acid evolutionary dynamics, which offers a baseline for evaluating observed amino acid patterns.

### 2. Mutational bias accounts for the predominance of amino acid substitutions throughout the COVID-19 pandemic

To quantitatively estimate the influence of mutational bias on the spectrum of amino acid substitutions across varying mutation-selection regimes, we necessitated a dataset characterized by substantial effective population sizes and elevated mutation rates. The exceptional genomic surveillance of SARS-CoV-2 presents this distinctive possibility. After several years of the COVID-19 pandemic, it became possible to rigorously validate our predictions by analyzing empirical amino acid substitution patterns in SARS-CoV-2. Comparing the observed rates of 150 single-nucleotide amino acid substitutions with those anticipated under a nearly neutral scenario (Fig. 1C) revealed a robust positive correlation, indicated by a Spearman correlation coefficient of 0.79 (N=150, p=3*10^-33^) (Fig. 2A) and a coefficient of determination (R²) of 59% (Supplementary Table 1). A negative control, based on randomly generated 12-component spectra, showed a markedly poorer model fit (Supplementary Fig. 2).

**Figure 2.**
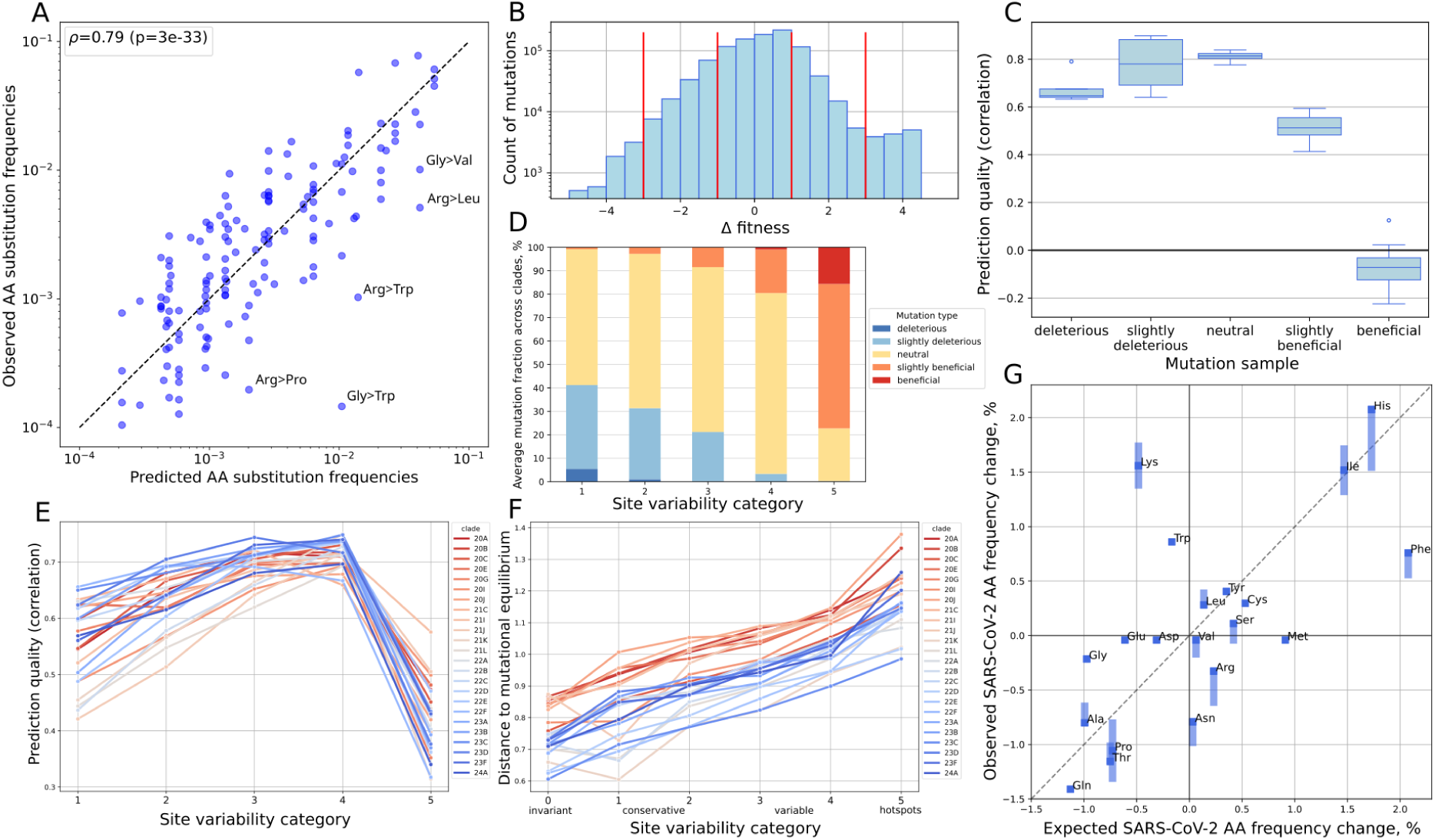
Observed SARS-CoV-2 amino acid substitutions during the COVID-19 pandemic in light of the nearly neutral model. (A) Observed versus predicted substitution frequencies for clade 20A, shown on a log-log scale. The black dashed line represents y=x. (B) The distribution of mutation fitness scores (Δfitness) for largest clade 21J (other clades have similar distributions). Vertical lines indicate thresholds for categorization. (C) Prediction quality of mutation samples, categorized by Δfitness. Spearman correlation estimated separately for 17 clades with at least 50,000 observed substitutions. Due to the lack of deleterious mutations in several clades, the sample correlation for this category was calculated using only five clades. (D) Distribution of average mutation fractions of different categories by Δfitness and by site variability. (E) Prediction quality of mutation samples, categorized by discrete gamma distribution. (F) Manhattan distance between observed in the proteome amino acid composition and expected mutational equilibrium composition across sites from different categories. Category 0 indicates invariant sites with no observed mutations during the pandemic. (G) Observed versus predicted amino acid frequency changes after 5 years of COVID-19. The black dashed line represents y=x. Mutational spectra are clade-specific in all analyses.

We further demonstrated that the model robustly predicts amino acid substitution frequencies across different clades and branch types (terminal or internal) (Supplementary Fig. 3), majority of proteins (Supplementary Fig. 4) and codon positions (Supplementary Fig. 5). Despite the overall relatively high prediction quality, substantial variation in model fit exists and warrants specific biological explanations. For example, the model fit was higher in the Omicron and later SARS-CoV-2 clades, fluctuated markedly across genes and tended to be higher on terminal compared to internal branches.

We hypothesised that fluctuations in model fit could be influenced by the strength of selection, the stronger it is – the stronger deviation from neutral amino acid substitution frequencies. The dense phylogenetic tree of SARS-CoV-2 recently enabled the assignment of fitness scores (Δfitness) to each amino acid substitution ^22^, providing an opportunity to test which substitutions align more closely with the nearly neutral scenario. By dividing all amino-acid substitutions into five categories, based on Δfitness (deleterious, slightly deleterious, effectively neutral, slightly beneficial, and beneficial) (Fig. 2B), we observed that neutral and slightly deleterious substitutions align very close to the model despite deleterious and (slightly) beneficial ones (Fig. 2C and Supplementary Fig. 6). This implies that the mutational spectrum is significantly distorted by selection; notably, positive selection appears to bias the spectrum more substantially than negative selection. Indeed, highly deleterious mutations remain unobservable in our dataset because viral genomes carrying such lethal variants are not viable and therefore cannot be sequenced.

A comparable pattern emerged when we analyzed substitution frequencies at the site level using gamma distribution to handle rate heterogeneity among sites (Supplementary Fig. 7). Sites under strong selective pressure—including conservative sites (categories 1–2), which are subject to purifying selection and enriched for deleterious mutations, and mutational hotspots (category 5), characterized by a high probability of beneficial mutations (Fig. 2D and Supplementary Fig. 8)—showed worse model fit than nearly neutral sites (Fig. 2E). This demonstrates that our model better captures the evolution of effectively neutral sites compared to either constrained sites or mutational hotspots under positive selection.

Next, we quantified the distance of the amino acid composition of SARS-CoV-2 proteome from the mutational equilibrium for different site categories. The amino acid composition of variable sites (categories 3–5) was significantly farther from the predicted equilibrium than that of conserved sites (categories 0–2) (Fig. 2F). This pattern is consistent with a nearly neutral evolutionary scenario when variable sites, being farther from equilibrium, are rapidly driven toward the mutational attractor (higher goddess of fit on Fig. 2E); conversely, conserved sites are already near equilibrium because purifying selection overrides the mutational forces. This is supported by the observed increase in GC fraction at variable sites (Supplementary Fig. 9). Alternatively, the large distance from equilibrium, particularly for mutational hotspots (category 5), may also result from positive selection, where beneficial mutations that deviate from the nearly neutral mutational spectrum shift the amino acid content away from the predicted equilibrium (Supplementary Note 1).

Finally, the overall evolution of amino acid composition was assessed for its consistency with the nearly neutral model. Proteome-wide amino acid frequency changes observed over five years of the COVID-19 pandemic were compared to predictions generated by Markow simulations using the Omicron mutation spectrum (clade 24A, see Methods). This analysis yielded a significant positive correlation (Pearson r = 0.646, p = 0.003, N = 20) between the observed and predicted changes (Fig. 2G). A consistent pattern was also evident in the empirical distribution of clade-specific amino acid mutations, as shown in the “empirical butterfly plot” (Supplementary Fig. 10).

The nearly neutral model, based solely on the mutational spectrum, explains at least 50% of observed amino acid substitutions in SARS-CoV-2 proteome and captures the amino acid composition change during the pandemic. Critically, the model’s predictive power correlates inversely with selection strength, demonstrating that deviations from the neutral expectation are a measurable proxy for purifying and positive selection pressures across different site categories.

### 3. Distinct mutational biases in +ssRNA, –ssRNA, and dsRNA viruses drive divergent proteome-wide evolutionary trajectories

To broaden our investigation beyond SARS-CoV-2, we examined additional viral species with distinct mutational spectra and assessed whether the observed amino acid divergence patterns correspond with their established mutational patterns. The accessibility of nucleotide sequences for various viruses, along with the automated mutation spectrum reconstruction methods (NeMu-pipeline) ^25^, enabled us to reconstruct mutational spectra for positive-sense transcripts of 32 viral species across 17 families, encompassing +ssRNA, −ssRNA, and dsRNA viruses (Supplementary Table 2). This collection was further validated (Supplementary Table 3) and extended with previously published spectra for eight additional viral species ^15^. The resulting dataset contains 35 spectra representing 34 species, including distinct profiles for the early (clade 20A) and late (clade 22C) SARS-CoV-2 lineages.

Species-specific mutational spectra reveal significant variance, particularly regarding the frequencies of four unique transitions (Fig. 3A, Supplementary Fig. 11). Among the families examined, Coronaviridae (Cov) displayed the most pronounced signature, marked by C>U and G>U biases. This is probably due to the nsp14-ExoN exoribonuclease domain, which corrects polymerase errors but is unable to repair damage caused by host immunological responses and inflammation ^26,27^. We also discerned distinct group-specific patterns, namely G>A (C>U in the genomic RNA strand) for –ssRNA viruses, and U>C for +ssRNA viruses (excluding coronavirids) (Fig. 3B). These findings align with other reports ^13,15^. The disparity between groups decreased after implementing complementary transformation on the spectra of –ssRNA viruses (Supplementary Fig. 12), indicating a universal mutational bias in RNA viruses, with C>U as the predominant substitution, which affects protein evolution in relation to genome sense.

**Figure 3.**
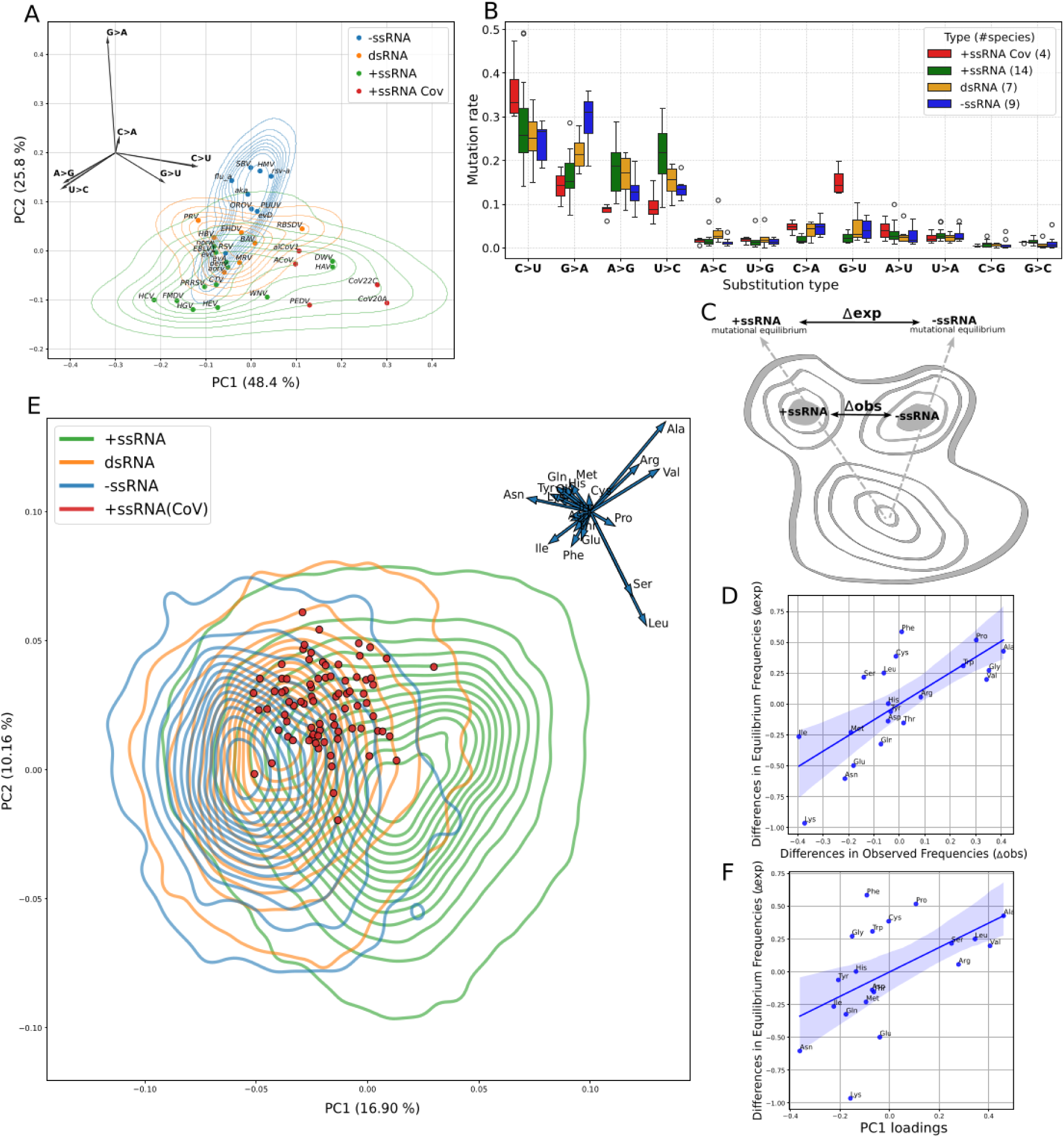
Impact of group-specific mutational spectra on RNA virus protein evolution. (A) Principal component analysis of viral mutational spectra; biplot origin adjusted to (-0.3, 0.2). (B) Group-specific mutational spectra distributions. (C) Scheme of divergent evolutionary trajectories toward mutational equilibrium for +ssRNA and –ssRNA viruses. (D) Correlation of predicted versus observed amino-acid compositional differences between +ssRNA and –ssRNA viruses, derived from aggregate spectra. Prior to correlation and subtraction, CLR-transformation was performed on the amino acid composition vectors. (E) Principal component analysis of RdRp amino acid content across 77,208 viral species; biplot origin set at (0.1, 0.1). (F) Relationship between macroevolutionary PC1 loadings from E and the predicted compositional shifts from D. “Cov” in the legends refers to coronavirids.

Given the distinct mutational spectra of +ssRNA (including coronavirids) and −ssRNA viruses, while dsRNA viruses typically have intermediate spectra (Fig. 3B), we focused on these two groups and compared the observed and expected differences in their average proteome-wide amino acid compositions according to the scheme in Figure 3C. The expected and observed differences are positively correlated (Pearson r=0.73, p=3*10^-4^, n=20) (Fig. 3D), demonstrating that group-specific mutational biases have shaped protein evolutionary trajectories regardless of selection. For example, the prominent G>A substitution bias in −ssRNA viruses suggests that the triplets for Ile, Asn, and Lys would be in excess in the plus-sense transcripts of −ssRNA viruses and in the relative deficit of +ssRNA viruses (Supplementary Fig. 13). Consistent with this prediction, Ile, Asn, and Lys are indeed in deficit in +ssRNA versus −ssRNA viruses in both the predicted and observed scenarios (Fig. 3D). The robustness of these patterns is further amplified by excluding viral lineages (*Coronaviridae)* where amino acid composition deviates markedly from equilibrium (Supplementary Fig. 14)—likely due to intense selection or recent shifts in the mutational spectrum—which yielded a more pronounced correlation (Pearson r=0.86, p=1.4*10^-6^, n=20) (Supplementary Fig. 15). This finding holds for species-level pairwise comparisons, with the relationship becoming more pronounced between virus pairs characterized by distinct mutational spectra and, consequently, more divergent evolutionary trajectories (Supplementary Fig. 16).

Finally, we investigated whether the influence of the mutational bias could also be discerned at the macroevolutionary scale. To address this, we concentrated on the RNA-dependent RNA polymerase (RdRp), the most conserved gene across all RNA viruses, for which amino acid sequences were available for 77,000 species ^28^ (Supplementary Fig. 17). Comparison of the amino acid composition of RdRp sequences across distinct virus groups demonstrated that, while these groups exhibited overlap, they presented group-specific compositional biases (Fig. 3E). Five amino acids (Asn, Ala, Val, Ser, Arg, and Leu) were found to significantly influence group separation, i.e. having strong loadings on the first principal component (PC1) (Supplementary Fig. 18). Based on nucleotide mutational bias, Asn is anticipated to occur at a high frequency in –ssRNA viruses, whereas Ala, Val, Ser, and Arg are specific to +ssRNA viruses (Supplementary Fig. 13). We also examined the relationship between the observed divergence in amino acid composition (PC1 loadings on Fig. 3E) and the predicted divergence driven by group-specific mutational spectra in +ssRNA and –ssRNA viruses. A significant positive correlation was observed (Fig. 3F) (Pearson r=0.53, p=0.02, n=20), demonstrating that nucleotide mutational bias is a principal determinant of the observed macroevolutionary differences in amino acid composition between these viral groups.

## DISCUSSION

Applying a nearly neutral framework to mutational spectra across diverse viral species, we find that intrinsic mutational biases explain a substantial fraction of observed amino acid substitutions, including more than half of substitutions in SARS-CoV-2 across site categories. This provides a quantitative baseline for separating mutation-driven trends from deviations more likely to reflect selection. At a broader evolutionary scale, group-specific mutational spectra are also associated with divergent proteome-wide trajectories in RdRp sequences from more than 77,208 viruses. Although RNA viruses provide an especially tractable system because of their large effective population sizes and high mutation rates, these results suggest that taxon-specific mutational bias can make a persistent contribution to protein evolution.

These findings support a view in which protein evolution is shaped jointly by mutation and selection, rather than by selection acting on an unbiased supply of variants ^29,30^. In this framework, mutational spectra influence which substitutions arise most often and therefore which adaptive paths are most accessible. Our results are thus consistent with mutation-biased adaptation, in which the mutation spectrum affects not only neutral change but also the order in which beneficial variants appear and the trajectories available to evolving populations ^3,7,31,32^. Selection still determines which variants persist, but mutational bias can channel the set of variants on which selection acts ^5,6,33^. Hence, considering the specificity of mutational spectra across diverse species and lineages ^13,15,34–36^, the evolutionary trajectories of each species on the tree of life are constrained by these spectra, in addition to the selection.

The relationship between mutational bias and selection is likely to depend on whether the dominant mutational tendencies of a lineage align with, or oppose, the locally favoured amino acid changes. When these forces act in the same direction, mutational bias may accelerate evolutionary change by increasing the supply of selectively tolerated variants. When they act in opposition, the same bias may instead constrain adaptation by making favourable substitutions less accessible. This view helps explain why the predictive contribution of mutational spectra can remain substantial across site categories while still varying with functional constraint, and it emphasizes that mutation bias should be understood not as an alternative to selection, but as a determinant of the variation on which selection acts.

This framework may have practical implications for evolutionary forecasting and engineering. In viral surveillance, estimates of mutational bias could help identify sites or regions whose future change is more or less constrained by the local supply of mutations. In biotechnology, deliberate shift of mutational spectra, for example through altered repair pathways ^8,9,37,38^, chemical mutagenesis ^39^, or engineered polymerases with distinct error profiles ^40–43^, may provide a way to bias evolving populations towards different regions of sequence space and probably extend it. These possibilities remain prospective, but they follow naturally from the observation that mutational input can shape the direction as well as the rate of protein evolution.

Several limitations should temper interpretation. Mutational spectra are not fixed and may change over time, across hosts, or with shifts in polymerase fidelity ^13,15^. Moreover, spectrum may be positively selected through adaptation to specific niches ^8,9^ (Supplementary Note 2). In addition, our use of simplified 12-component spectra does not capture adjacent-nucleotide context, e.g. CpG, which can materially affect mutation probabilities ^13,44,45^. Predictive power should therefore be interpreted as an indirect measure of the joint effects of mutation and selection, rather than as a direct estimate of selection itself. Extending this framework to context-dependent spectra and time-resolved mutational processes should improve both mechanistic resolution and predictive accuracy. These limitations likely explain part of the remaining unexplained variance and point to clear extensions, including context-dependent spectra and time-resolved models of mutational change.

More broadly, the results support a view of protein evolution in which mutation bias and selection act jointly rather than independently. Selection determines which variants persist, but mutational spectra influence which variants arise most often and therefore which evolutionary paths are most accessible. Quantifying this mutational component makes it possible to move beyond descriptive accounts of sequence change towards a more predictive framework, both for short-term viral evolution and for long-term divergence across taxa.

## MATERIALS AND METHODS

### Dataset of SARS-CoV-2 mutations

Information regarding SARS-CoV-2 mutations and their corresponding mutational spectra, derived through phylogenetic analysis, was sourced from Bloom and Neher ^22^. This comprehensive dataset (filtration parameters: subset=all, exclude=false) comprised 5,635,999 genomic mutations, encompassing both synonymous and missense types. Additionally, it provided fitness coefficient values for every mutation across 24 specific virus clades and lineages, along with 12-component spectra tailored to each clade.

### Dataset of RNA-viruses mutational spectra

Mutational spectra for representative 32 RNA viruses were reconstructed using the NeMu-pipeline^25^. Inclusion criteria were limited to viral species with more than 500 nucleotide sequences available in GenBank, as detailed in Supplementary Table 4. Sequence abundance data were retrieved from the Wikidata-based Taxonium project phylogenetic tree ^46^. The analytical pipeline focused on highly sequenced protein-coding regions: for viruses possessing a single polyprotein, complete genome sequences were utilized, whereas for other viruses, analysis was restricted to coding sequences of at least 3 kb. Briefly, the NeMu-pipeline performed orthologous sequence retrieval from GenBank via tblastn, reconstructed the associated phylogenetic trees, and inferred single-nucleotide substitution probabilities from the tree branches. To establish nearly neutral mutational spectra, only synonymous mutations were utilized in the derivation.

The mutational spectra for RSV-A, RSV-B, Enteroviruses (D68 and A71), West Nile virus, influenza A and B, and dengue viruses were obtained from the Supplementary materials provided in ^15^. To validate our computational reconstructions, we performed a pairwise cosine similarity analysis against these established profiles, which demonstrated exceptional concordance with values exceeding 0.99 (Supplementary Table 3). By integrating reference spectra of SARS-CoV-2 (clades 20A and 22C) and RSV-A with our reconstructions for 32 species, we compiled a comprehensive dataset consisting of 35 unique viral spectra across diverse lineages (Supplementary Table 2).

### Dataset of viral RdRp sequences

We acquired 77,208 RNA virus RdRp sequences from the Trees and Alignments section of the Serratus database ^28^ (https://serratus.io/trees, accessed May 15, 2022). The viruses included in this dataset span 24 orders and 73 families (Supplementary Fig. 17). Across the viral dataset, the median length of the sequences was 105 amino acids. Following the calculation of the amino acid content for each sequence we applied Principal Component Analysis (PCA) to the resulting dataset.

### Predicting the amino acid substitution frequencies based on mutational spectrum

We modeled amino acid substitution frequencies by integrating 12-component nucleotide mutational spectra µ, estimated from synonymous substitutions, with the standard genetic code. We calculated the transition probabilities for each codon to all accessible single-nucleotide variants based on substitution frequencies in the spectrum, assuming null rates for substitutions requiring multiple nucleotide changes following the approach in ^3,4,23,24^:

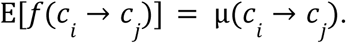

In such a way we derived a codon substitution matrix (64x64). By aggregating these frequencies, we constructed a matrix of relative amino acid substitution frequencies (20x20), with distinct entries for forward and reverse rates. This matrix contains 150 non-zero frequencies according to the standard genetic code.

### Estimating the mutational equilibrium frequencies

Equilibrium amino acid frequencies were calculated based on equilibrium codon frequencies aggregation. For this, the codon substitution matrix Q, obtained from the mutational spectrum as described above, was used. Equilibrium frequencies were calculated analytically as the real part of the principal eigenvector of matrix Q, corresponding to the zero eigenvalue. The approach is implemented in Python based on the scripts used to calculate equilibrium nucleotide frequencies in ^15^. Manhattan distance was used to compare observed and equilibrium amino acid frequencies.

By comparing empirical amino acid frequencies with predicted equilibrium values, we estimated "neutral trends" for frequency shifts within specific proteins under defined mutational spectra. This analysis allowed us to identify sets of amino acids categorized as "gainers"—those expected to increase in frequency under a nearly neutral evolutionary scenario—and "losers," whose proportions are projected to decline over time.

### Validation of the model of amino acid substitutions

For every mutation dataset, we derived expected amino acid substitution rates from the specific mutational spectrum and evaluated them against empirical patterns in a lineage-specific manner. To determine empirical frequencies for amino acid substitutions, mutations were analyzed across various genomic regions and specific site samples, with adjustment made to account for the baseline amino acid composition of those sites.

The robustness of our model was assessed using Spearman and Pearson correlation coefficients, the coefficient of determination (R^2^), and the weighted mean absolute percentage error (WAPE). To account for the inherent constraints of compositional data, we applied a Centered Log-Ratio (CLR) transformation^47^ to the substitution frequency vectors prior to calculating parametric statistical metrics using the following equation:

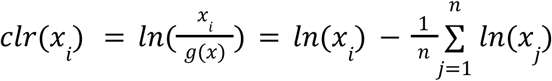

where *g(x)* is the geometric mean of the substitution frequency vector *x*.

To establish a negative control, we assessed the model’s robustness by substituting lineage-specific nearly neutral profiles with randomly generated mutational spectra. Utilizing a uniform distribution to sample nucleotide substitution rates across 20 independent replicates, we recalculated the predictive metrics to quantify the expected degradation in model fit (Supplementary Fig. 2).

### Discrete modeling of site-specific variability

We employed a zero-inflated, composite discretization framework to analyze rate heterogeneity across different sites within each dataset (Supplementary Fig. 7). Missense mutation frequencies for individual sites were normalized against the global average to determine relative evolutionary rates. Sites lacking any empirical substitutions were classified as ‘invariant’ (category 0). To isolate extreme outliers, we established a ‘hotspot’ group (category 5) for sites exceeding the 99th percentile of the variability distribution. The remaining site-specific rates were approximated using a Gamma distribution—optimized through maximum likelihood estimation—and subsequently partitioned into four equal-probability categories. This hierarchical discretization prevents invariant positions and hyper-mutable outliers from distorting the underlying distribution’s shape parameter.

### Simulating proteome-wide amino acid divergence during the COVID-19 pandemic

To evaluate evolutionary shifts within the SARS-CoV-2 proteome, we developed a nearly neutral null model. Amino acid transition probabilities were derived from an instantaneous rate matrix Q, parameterized by the Omicron mutational spectrum (clade 24A). The initial state vector (π_0_) was defined by the ancestral Wuhan strain sequences.

Expected evolutionary trajectories were modeled via a continuous-time Markov chain (CTMC). We calculated the time-resolved vector of amino acid frequencies (π_t_) using the matrix exponential function, numerically integrated across discrete steps according to the following expression:

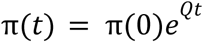

where π represents the frequency vector and *t* denotes the temporal increment. Simulation was terminated when the modeled genetic distance reached parity with the empirical divergence observed in 2025 lineages relative to the reference. This allowed for a direct comparison between observed proteomic shifts and a nearly neutral scenario driven exclusively by mutational bias.

Empirical frequency fluctuations (Δf_aa_) were quantified as normalized deviations from ancestral Wuhan strain values across 59,361 genomes sampled between January 1 and May 22, 2025 (GRA lineages), utilizing the following equation:

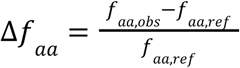

Pearson correlation was used to assess the correspondence between empirical data and simulated neutral trajectories for each genome. The mean correlation coefficient served as a quantitative metric for the predictive power of the mutational spectrum in shaping the SARS-CoV-2 proteome-wide amino acid landscape.

### Comparison of observed and expected amino acid content differences

Utilizing lineage-specific mutational spectra, we derived the expected equilibrium amino acid frequencies for individual viral species and calculated group-level aggregates for −ssRNA and +ssRNA viruses. Concurrently, empirical amino acid compositions were determined for each virus and averaged within their respective groups. To address the inherent constraints of compositional data, we applied a Centered Log-Ratio (CLR) transformation^47^ to the resulting vectors. The predicted evolutionary divergence was quantified by subtracting −ssRNA from +ssRNA equilibrium frequencies, with observed differences calculated through a parallel subtraction of empirical group frequencies. The statistical correspondence between these predicted and observed shifts was evaluated using Pearson correlation, following a Shapiro-Wilk normality assessment. This comparative framework was additionally extended to analyze pairwise species interactions beyond aggregate group trends.

## DATA AVAILABILITY

Dataset of 32 viral mutational spectra, estimated using NeMu-pipeline, and all of our code used to obtain all figures are available at https://github.com/mitoclub/RNAViralMutSpec. Dataset of 4,132,247 mutations and clade-specific spectra of SARS-CoV-2 reconstructed using a phylogenetic approach were derived from materials of ^22^ and are available at https://github.com/jbloomlab/SARS2-mut-fitness/tree/main/results. Dataset of 13 viral mutational spectra for eight species was derived from materials of ^15^ and is available at https://github.com/jbloomlab/SARS2-mut-spectrum/tree/main/results/other_virus_spectra.

## ACKNOWLEDGMENTS

We thank Emma Penfrat, Melissa Franco, Zoe Fleischmann, Arina Trufanova, Thomas Junier for participation in the data analysis on the early stages of this research; and Sergey Oreshkov, Stepan Denisov, Georgii A Bazykin and Evgenii Tretiakov for fruitful discussions.

Conflict of interest statement. None declared.

## FUNDING

This work was supported by Russian Science Foundation, Grant № 21-75-20143.

## AUTHOR CONTRIBUTIONS

K.P. and K.G. designed research; B.E., A.V., V.S. and E.P. performed research and analyzed data; K.P., K.G., J.F., K.K., A.A., B.E., V.Y. and V.T. discussed the results; K.P. and B.E. wrote the paper. K.G. and A.A. proofread the paper.

## Notes

### Competing Interest Statement

The authors have declared no competing interest.

