## Supplementary Figures for "Mutational bias shapes protein evolution across RNA viruses"

Efimenko B. et al.

**Supplementary materials**

Supplementary notes 2
Note 1. The hypothesis of biased amino acid content from mutational equilibrium 2
Note 2. The mutational spectrum can be pre-adaptive 2
Supplementary figures 5
Figure 1. Mutational spectrum of synonymous mutations for different site samples. 5
Figure 2. Quality estimation of the model of amino acid substitutions on random spectra for SARS-CoV-2 clades. 6
Figure 3. Quality estimation of the model on different SARS-CoV-2 clades and tree branches subsets. 6
Figure 4. Quality of amino acid substitution frequencies prediction on SARS-CoV-2 proteins for early and late clades. 7
Figure 5. Quality estimation of the model on mutation samples at different codon positions in SARS-CoV-2 genes. 8
Figure 6. The model quality on mutation samples with different delta fitnesses for SARS-CoV-2 clades. 8
Figure 7. Site gamma-categorization based on variability for SARS-CoV-2 genome. 9
Figure 8. Distribution of average mutation counts of different categories by Δfitness and by site variability 10
Figure 9. GC content of codon sites in different site categories according to variability. 10
Figure 10. SARS-CoV-2 amino acid composition changes during COVID-19 pandemic. 11
Figure 11. Viral mutational spectra based on synonymous mutations computed using NeMu-pipeline. 12
Figure 12. Mutational spectra of mutated strand of RNA-viruses. 13
Figure 13. Amino acid mutational equilibrium frequencies distributions for 4 groups of viruses. 14
Figure 14. Distance to equilibrium amino acid composition for viruses in our dataset. 15
Figure 15. Comparison of observed versus expected differences in amino acid composition between +ssRNA and –ssRNA viruses excluding distant from equilibrium viruses. 16
Figure 16. Pairwise comparison of observed and expected differences in amino acid content among 34 viral species. 17
Figure 17. Comprehensive overview of RNA virus species RdRp sequence dataset. 18
Figure 18. The amino acids loadings in the first and second principal components of RdRp amino acid composition PCA-analysis. 19

### Supplementary notes

#### Note 1. The hypothesis of biased amino acid content from mutational equilibrium

We asked how far the amino acid composition of different sites lies from the expected neutral amino acid equilibrium. There are 4 scenarios, depending on the main driver of compositional change – mutagenesis or selection, or a coupling of both:

1) The mutagenesis-driven scenario assumes that rapidly evolving, effectively neutral positions (categories 3–4) show high goodness of fit of the model because they are far from equilibrium and mutational forces drive them quickly toward the mutational attractor; in contrast, conserved categories (1–2) are already close to equilibrium, and mutational forces are weaker relative to purifying selection there.

2) The negative selection-driven scenario assumes that conserved categories evolve under selection that acts independently of the mutational attractor and shapes variability separately from mutagenesis, thereby keeping them farther from the mutational equilibrium, whereas nearly neutral sites are pushed closer to equilibrium by the mutational process due to weaker selection.

3) The positive selection-driven scenario assumes that positive mutations that oppose the mutational bias move out the amino acid content from equilibrium.

4) The biased-to-adaptive mutational architecture scenario assumes that the mutational spectrum itself is an evolved, first-order driver of adaptation. In this view, mutagenesis and selection are fundamentally aligned: the mutational attractor is structurally biased to "undersample" mutations at sensitive conserved sites (preserving functional integrity) while providing complexity-preserving adaptive paths for variable sites.

Interestingly, our results are consistent with the mutagenesis-driven scenario, which can point toward an biased-to-adaptive mutational architecture: the amino acid composition of variable sites is farther from equilibrium than that of conserved sites (Fig. 2). Although more complex scenarios involving interactions between selection and mutagenesis (as formalized in scenario 4) are possible, this pattern provides a simple rule of thumb—conserved sequences can be conserved not only because of strong negative selection but also because they are evolved mutational cold spots (e.g., where the biased spectrum limits mutational input), i.e., they already contain many “gainers” and few “losers” and therefore mutate slowly, whereas variable sites are variable precisely because they are dominated by “losers” and evolve rapidly toward the “gainers” provided by the biased mutation spectrum.

#### Note 2. The mutational spectrum can be pre-adaptive

The gain and loss of the repair genes in bacterial phylogeny occur at a rate much higher than previously anticipated. This frequent "spectral flux" suggests that populations can rapidly modulate their mutational input in response to environmental shifts, effectively "choosing" a mutational spectrum that provides the highest probability of survival in a new niche. [doi:10.1073/pnas.2207355120] As a population resides in a constant environment for an extended period, it undergoes an "adaptive walk" toward a fitness optimum. During this process, natural selection efficiently identifies and fixes the most accessible beneficial mutations provided by the existing mutational bias. Consequently, the "pool" of beneficial mutations within that biased class becomes depleted. This leads to a situation where the remaining possible mutations in that class are largely neutral or deleterious, resulting in a "left-shifted" Distribution of Fitness Effects (DFE) for the ancestral bias. [doi:10.1371/journal.pbio.3003282] E. coli strains that reversed the ancestral transition bias (ΔmutY, ΔmutT) consistently exhibited right-shifted DFEs, characterized by a higher proportion of beneficial mutations and a lower deleterious load. Conversely, strains that reinforced the transition bias (ΔmutS) showed left-shifted DFEs, with significantly fewer beneficial outcomes. [doi:10.1371/journal.pbio.3003282] The identification of genes that normally increase the mutation rate—whose disruption results in an "antimutator" phenotype—provides a novel target for drug development. Antimutator strains, such as ΔnudJ in E. coli, show reduced mutation rates and shifted spectra that limit access to the fittest mutations. Future "anti-evolution" drug strategies could focus on inhibiting these pathways during antibiotic or chemotherapy treatment. By suppressing the cell's ability to undergo adaptive spectrum shifts (e.g., bias reversals), these drugs could prevent the emergence of resistance and keep the population trapped on a fitness plateau where its beneficial mutations have been depleted. [doi:10.1093/molbev/msaf182] Moreover, it was found that different mutational paths to the same level of antibiotic resistance—driven by different mutational biases—result in markedly different "collateral sensitivity profiles". For example, a population that acquires resistance through a transition-biased path may remain susceptible to a second antibiotic, while one that takes a transversion-biased path may develop cross-resistance. [doi:10.1038/s41467-026-74044-6] This implies that the adaptiveness of the mutational spectrum has far-reaching effects on the predictability and long-term viability of a population. The "direction" of evolution, dictated by the spectrum, determines the population's future adaptive potential and its vulnerability to subsequent environmental challenges. [doi:10.1098/rstb.2022.0055]

Human germline evolution has been shaped over millions of years to minimize the occurrence of mutations in sensitive, functional regions. Consequently, the germline spectrum "oversamples" certain classes of mutations and "undersamples" others. The mutations required to drive cancer—specific driver mutations in genes like TP53, PIK3CA, and KRAS—often reside in the classes that the germline has historically avoided. In Nonhypermutator (NHM) cancers, the mutational spectrum is anticorrelated with the germline likelihood. This means that the mutations most frequently produced by the tumor's altered spectrum are precisely those that were rarest in the germline. [doi:10.1093/molbev/msaf105]. NHM tumors, which maintain relatively low mutation rates compared to their hypermutator (HM) counterparts, show a conserved shift in their mutational spectrum across more than 20 different tissue types. [doi:10.1093/molbev/msaf105] This implies that the diversification of mutation rates and spectra is driven by developmental and evolutionary pressures. Paco Majic et al. suggest the hypothesis that the timing and location of mutational events are far from random; they are structured to balance the need for innovation with the preservation of functional integrity. [doi:10.1101/2025.01.16.633395]. A critical piece of evidence for the adaptiveness of the mutational spectrum in developmental processes is the discovery of "programmed" DNA breakage. Somatic mutations in the human brain accumulate in a non-random, structured manner. Specific signatures are enriched in regulatory regions involved in housekeeping and cell maintenance. This "functional mosaicism" suggests that the neuronal genome is managed as a dynamic landscape where programmed damage and biased repair balance long-term integrity with short-term plasticity. [doi:10.1038/s41586-021-03468-5; doi:10.1038/s41588-021-01001-y]

The adaptiveness of the mutational spectrum is equally evident in protein evolution, where the spectrum of adaptive substitutions reflects the underlying mutation bias and the biophysical constraints of protein folding. Statistical analysis of protein sequence adaptation across diverse species shows that the mutation spectrum has a proportional influence on the types of changes fixed in evolution. For a given beneficial amino acid change, the probability of its fixation is heavily weighted by the rate at which that specific mutation is generated. [doi:10.1073/pnas.2119720119] Moreover, the connectivity of the genetic code and the nature of amino acid substitutions suggest that the mutational spectrum is aligned with structural and functional preservation. Amino acids that are similar in molecular complexity are more likely to be connected by single-point mutations than would be expected by chance. This structural connectivity is adaptive because it ensures that most mutations lead to "complexity-preserving" substitutions, which are less likely to destabilize the protein fold. Therefore the evolution of the genetic code itself may have been constrained to maximize mutational accessibility between biochemically compatible amino acids, minimizing the energetic costs of new functional innovations. [doi:10.1093/gbe/evag012]

Thus, these evidences confirm that the mutational spectrum is a first-order driver of adaptation, determining the rate, direction, and predictability of evolution. In other words, the paradigm shift from "random mutation" to "adaptive mutational architectures" and this conceptual change  marks a significant advancement in our ability to decipher and direct the course of evolution.

### Supplementary figures

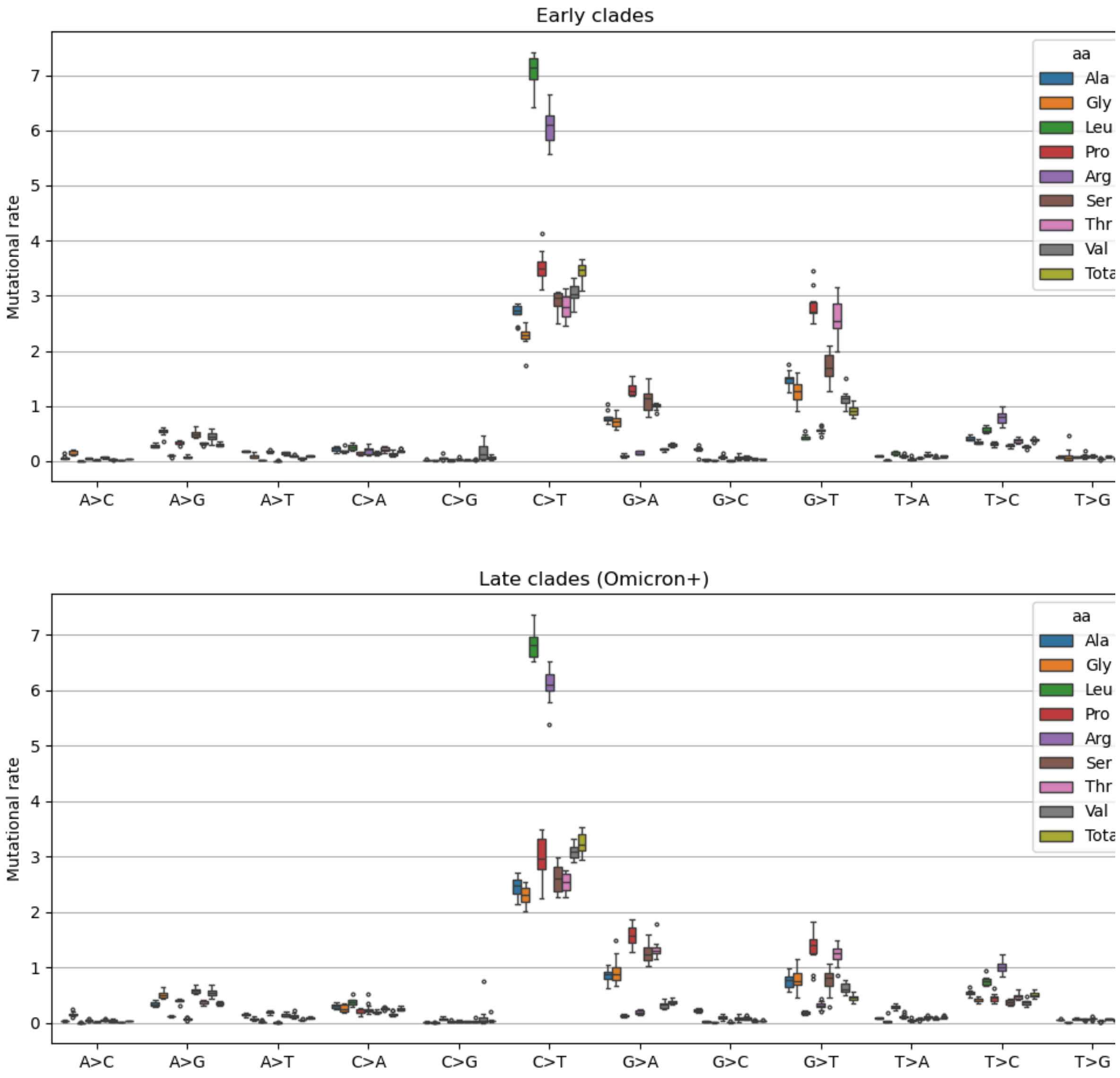


**Figure 1. Mutational spectrum of synonymous mutations for different site samples.** Each sample is based on fourfold sites that belong to codons, which encodes distinct amino acids (see legend). There are strict mutational biases for mutations in codons of Leu and Arg, but generally the total spectrum is highly similar to other spectra.

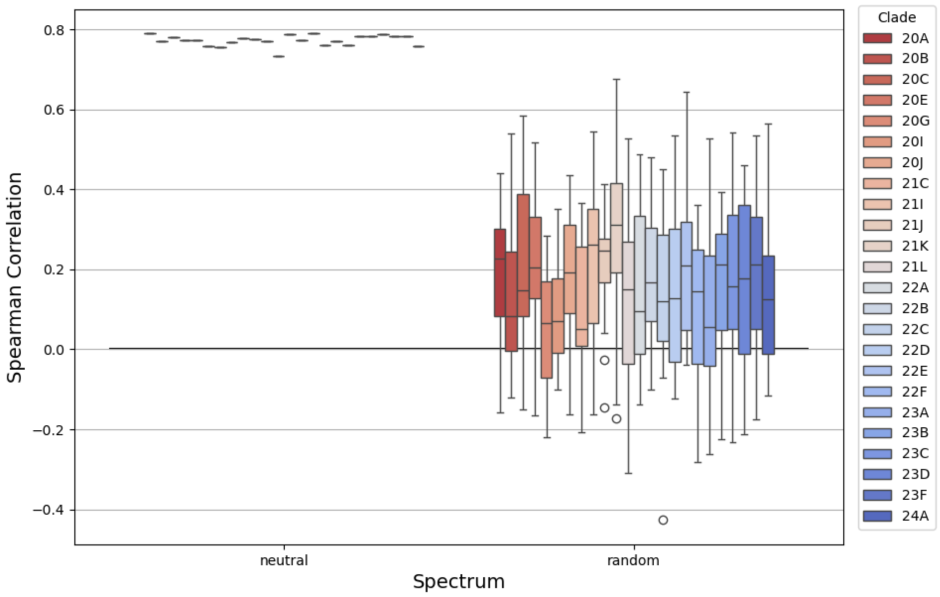


**Figure 2. Quality estimation of the model of amino acid substitutions on random spectra for SARS-CoV-2 clades.**

Used 24 clades. The "neutral spectrum" is the clade-specific spectrum based on synonymous mutations, while the random spectrum sampled 20 times from uniform distribution for each clade.

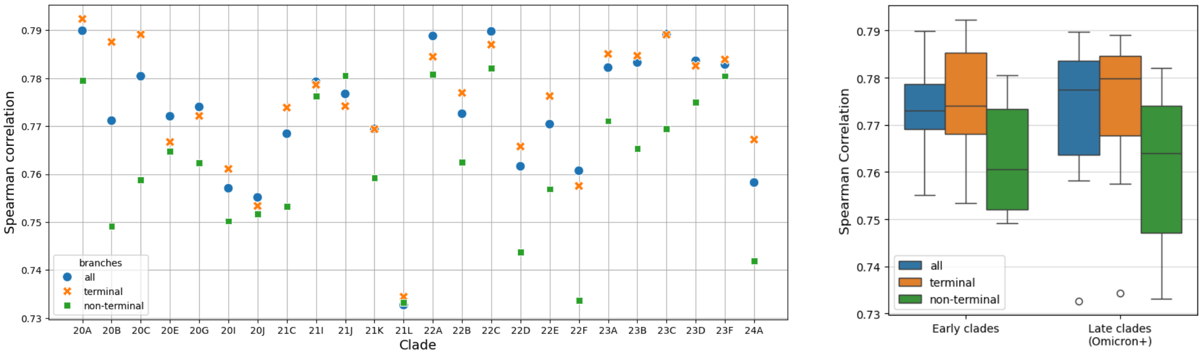


**Figure 3. Quality estimation of the model on different SARS-CoV-2 clades and tree branches subsets.**

Clade 21K is the first Omicron clade during the pandemic.

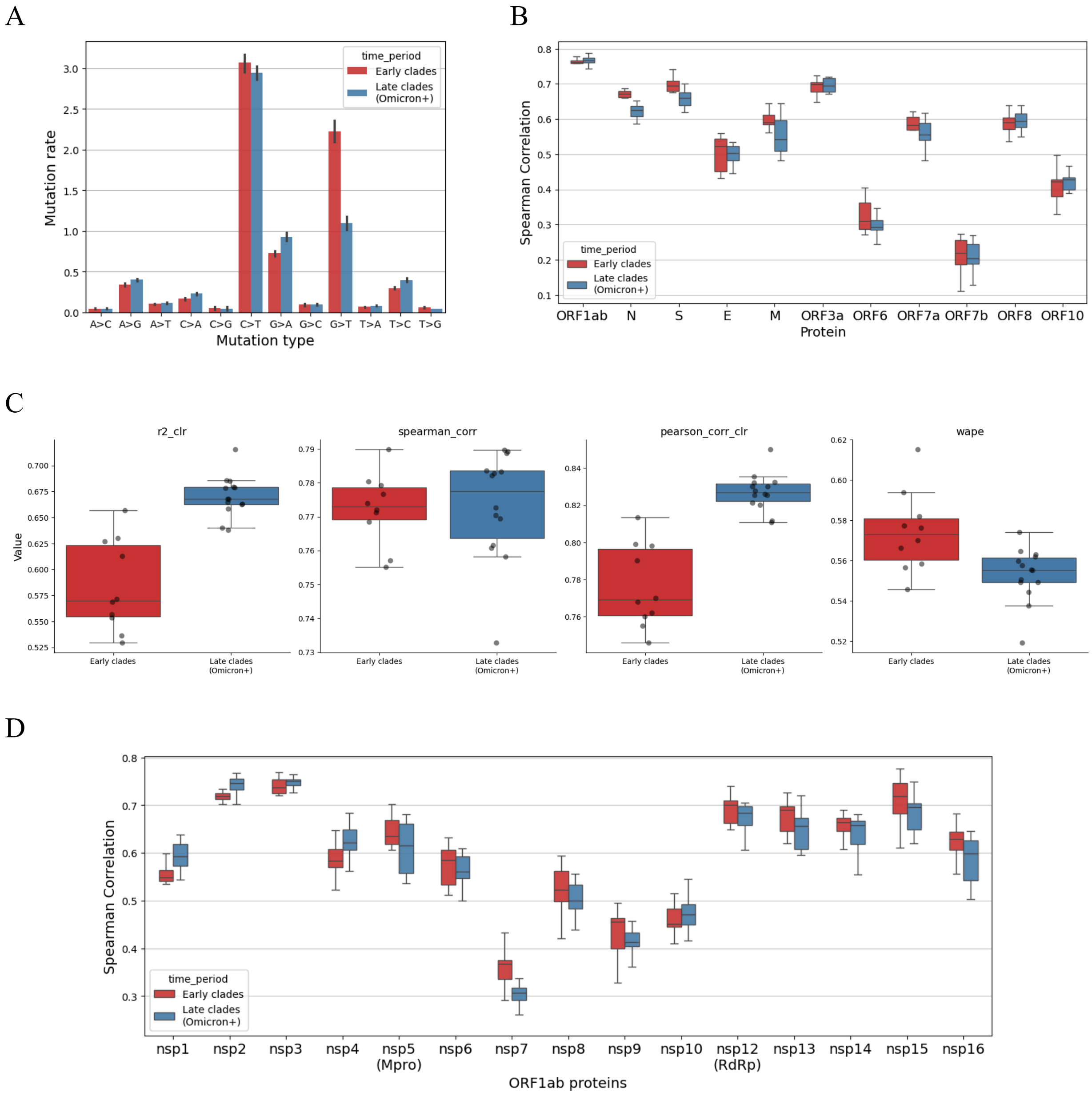


**Figure 4. Quality of amino acid substitution frequencies prediction on SARS-CoV-2 proteins for early and late clades.**

(A) Average mutational spectra for early and late (Omicron-specific) clades. (B) Model quality for SARS-CoV-2 genes. (C) Quality estimation using several metrics for early and late clades. "r2" is the coefficient of determination. "_clr" labels indicate that the CLR-transformation was applied to observed and expected substitution frequencies before the metrics were calculated.

(D) Model quality for proteins, encoded by the ORF1ab gene.

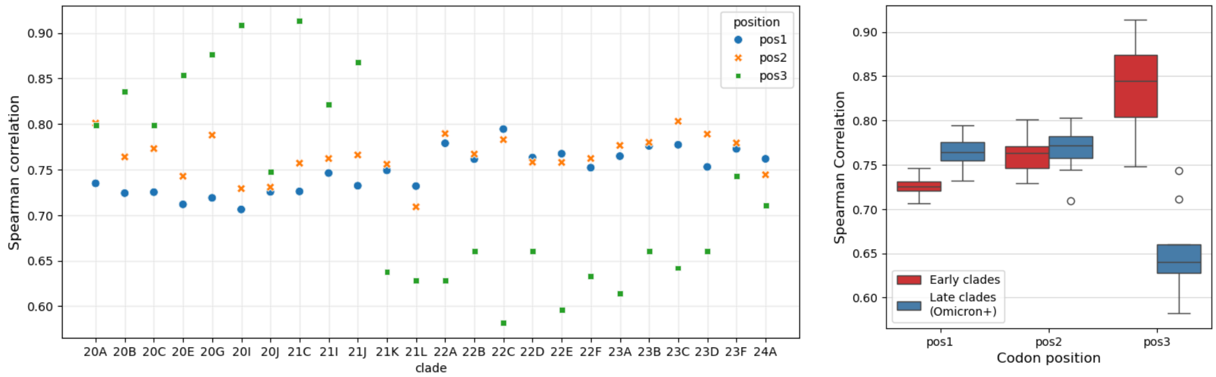


**Figure 5. Quality estimation of the model on mutation samples at different codon positions in SARS-CoV-2 genes.**

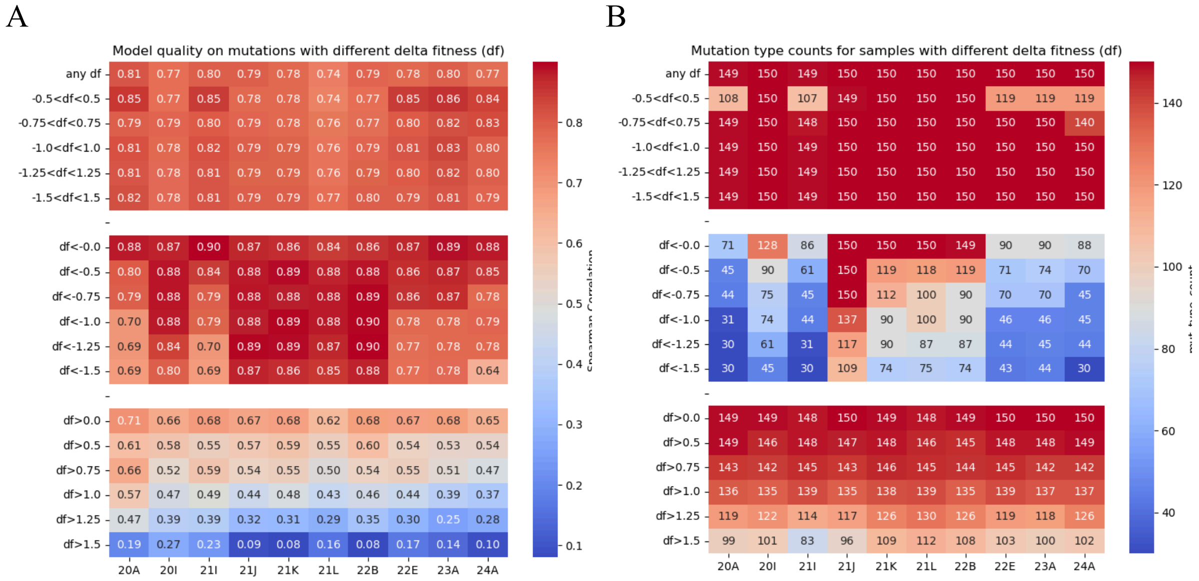


**Figure 6. The model quality on mutation samples with different delta fitnesses for SARS-CoV-2 clades.**

(A) Heatmap with Spearman correlation coefficient values for different samples of mutations. “df” is the delta fitness score of mutation. (B) Similar heatmap with substitution type counts. The maximum number of amino acid substitution types is 150 according to the standard genetic code.

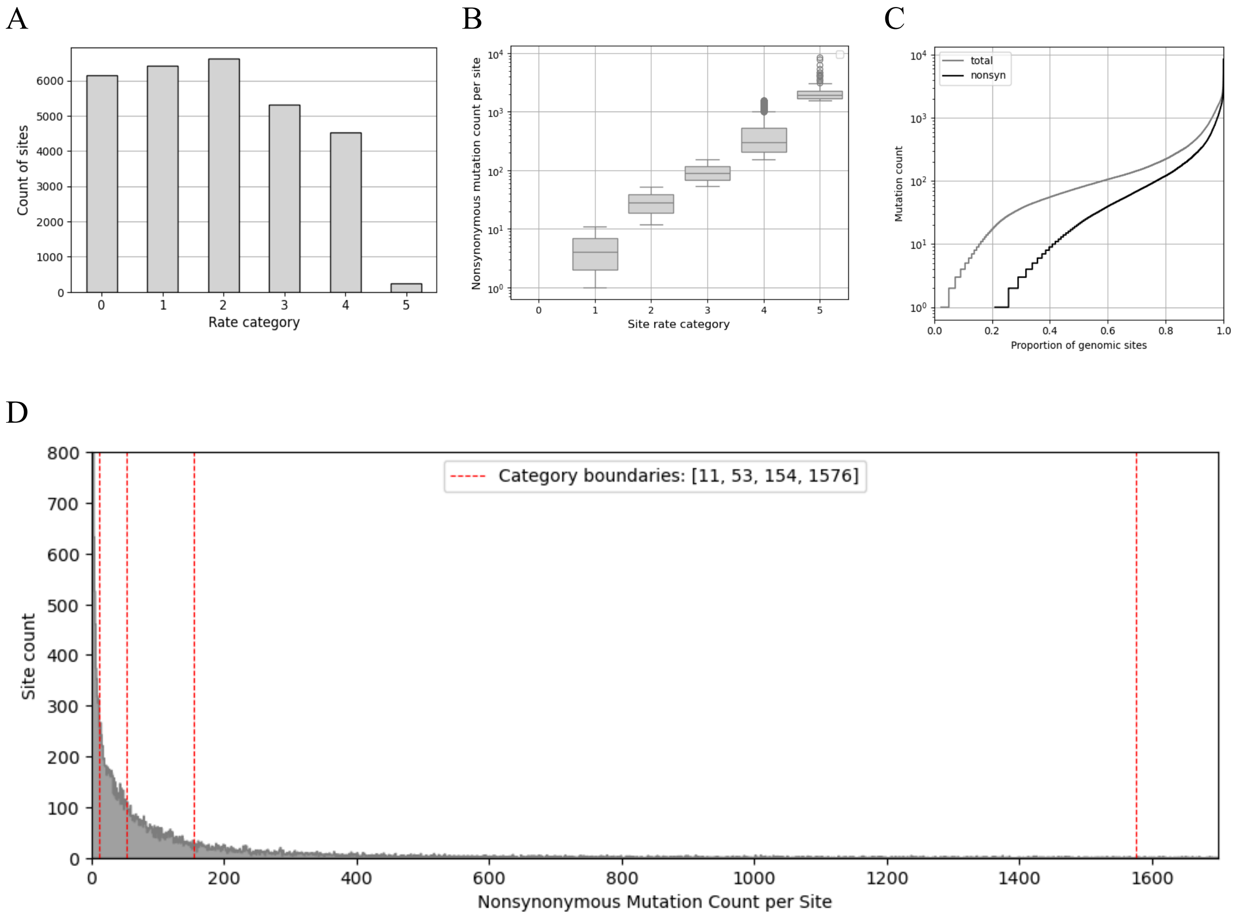


**Figure 7. Site gamma-categorization based on variability for SARS-CoV-2 genome.**
(A) Count of sites across different mutation rate categories, with variability increasing from category 0 (invariant sites) to 5 (mutational hotspots). (B) The distribution of missense mutation count per site in the gamma-categories. (C) Empirical cumulative distribution of missense mutations in the genome. (D) The distribution of missense mutation count per site. Categorization boundaries indicated by red dashed lines. The plot is limited in both axes.

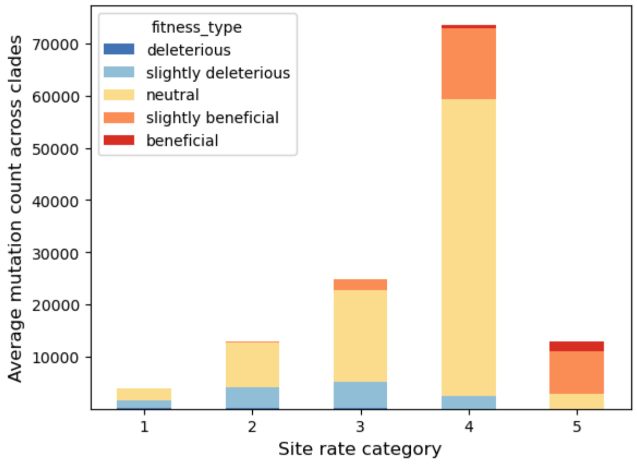


**Figure 8. Distribution of average mutation counts of different categories by Δfitness and by site variability**

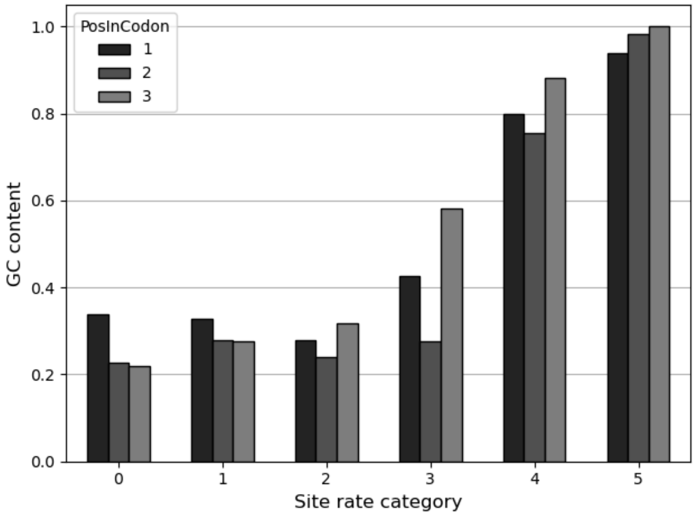


**Figure 9. GC content of codon sites in different site categories according to variability.**

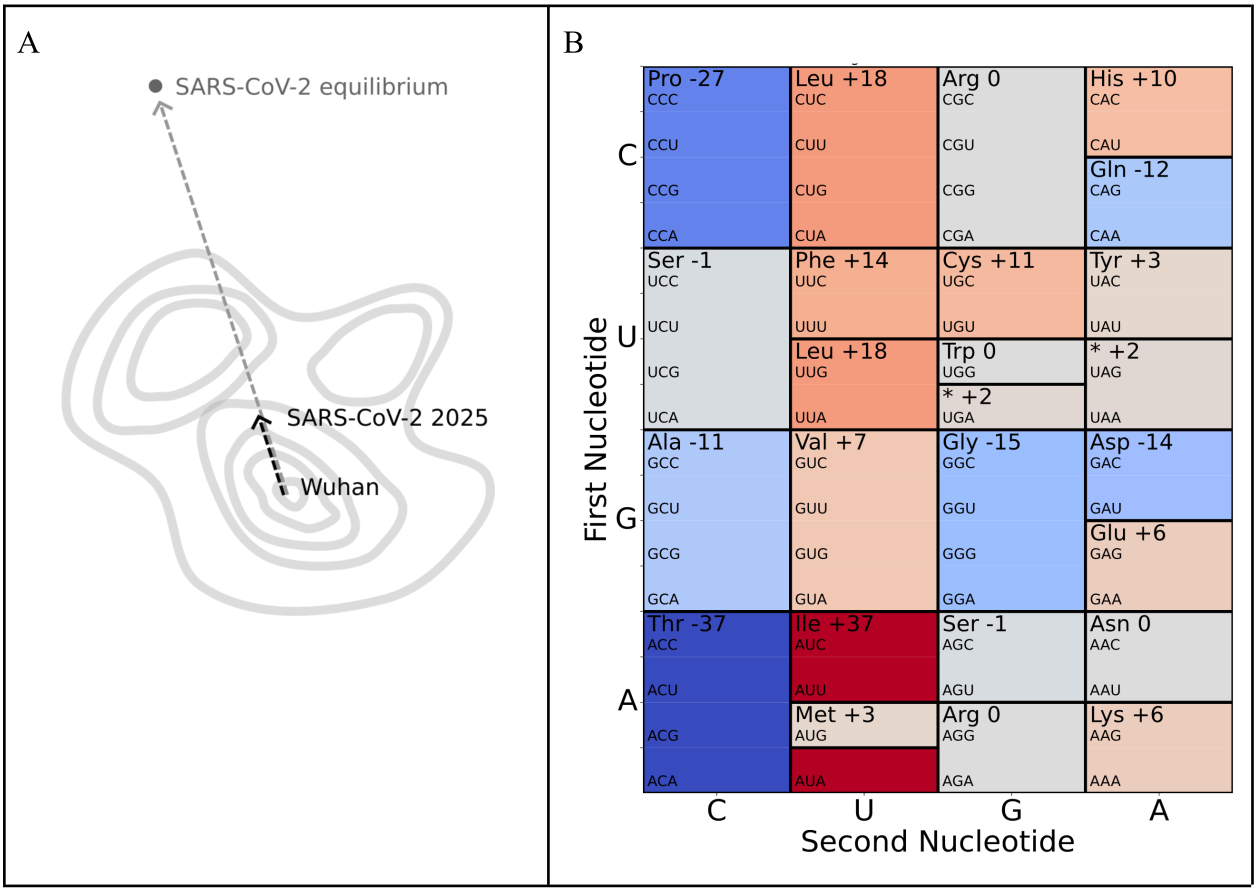


**Figure 10. SARS-CoV-2 amino acid composition changes during COVID-19 pandemic.** (A) Scheme of the virus proteome evolution at fitness landscape space. (B) Counts of distinct amino acid changes depicted on the standard genetic code table. Used only clade-specific amino acid substitutions that probably were positively selected in different clades and improved fitness of the virus (Table 2 in https://doi.org/10.1093/molbev/msad085). Notably, this empirical butterfly plot closely resembles the theoretical one (see Fig. 1B).

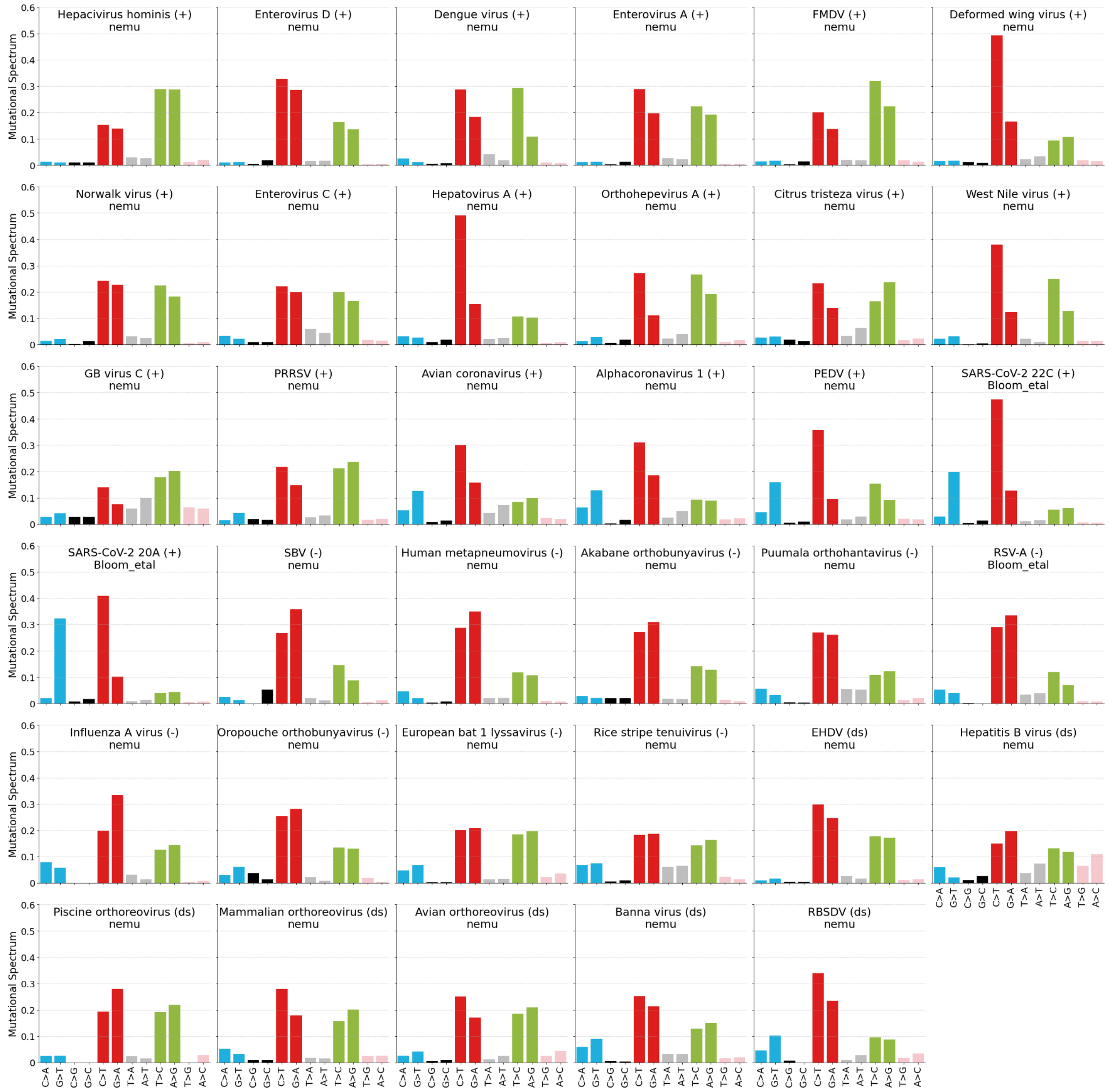


**Figure 11. Viral mutational spectra based on synonymous mutations computed using NeMu-pipeline.**

RSV-A and SARS-CoV-2 spectra are derived from https://doi.org/10.1093/molbev/msad085

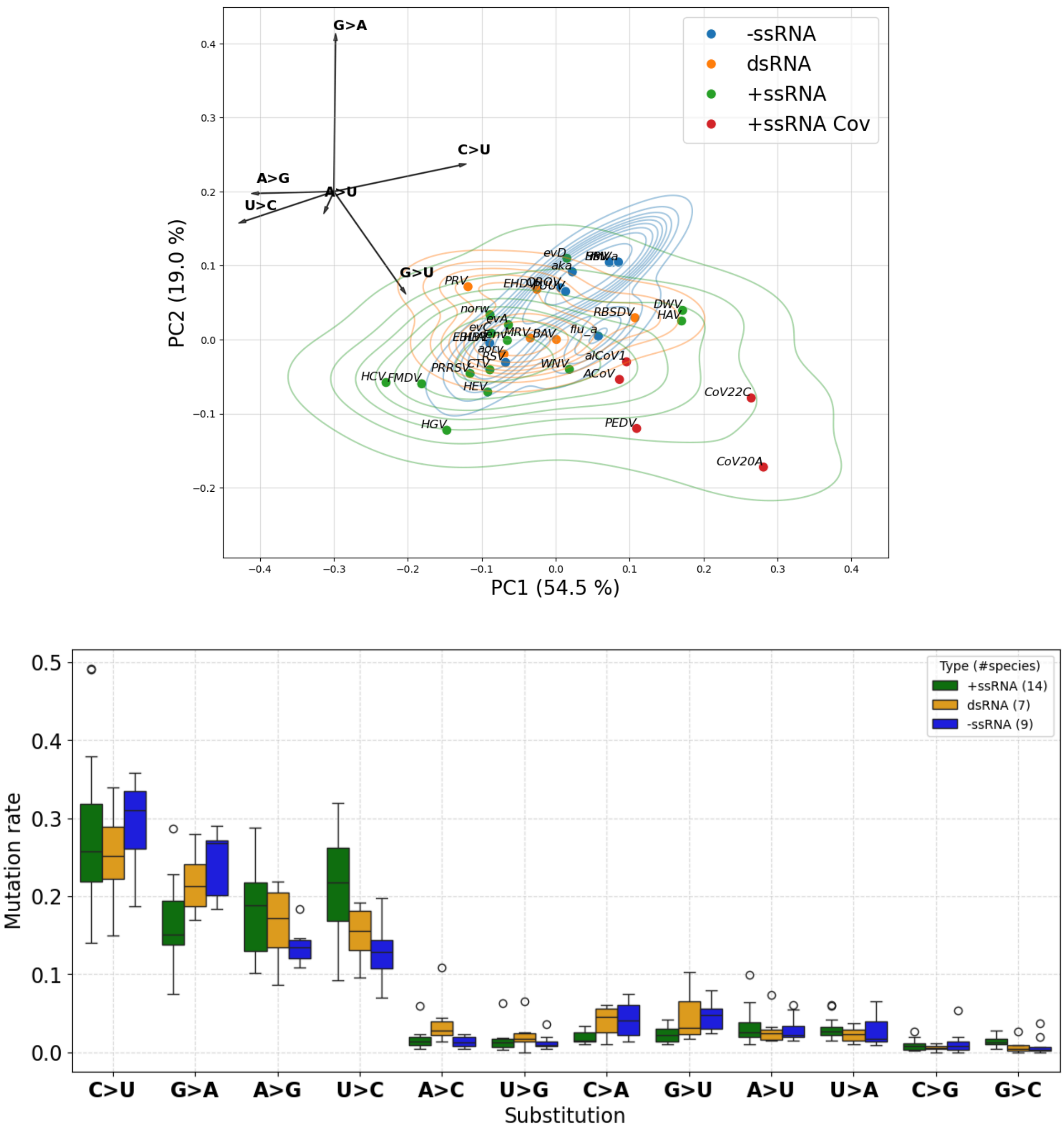


**Figure 12. Mutational spectra of mutated strand of RNA-viruses.**

PCA on upper panel and barplots on bottom panel.

To account for the higher concentration of negative-sense genomes in host cells and their presumed role as the primary site of mutation accumulation, we performed a complementary transformation on the spectra of positive-sense transcripts from –ssRNA species. In contrast, the spectra for +ssRNA and dsRNA viruses remained untransformed, consistent with the Figures 3A and 3B.

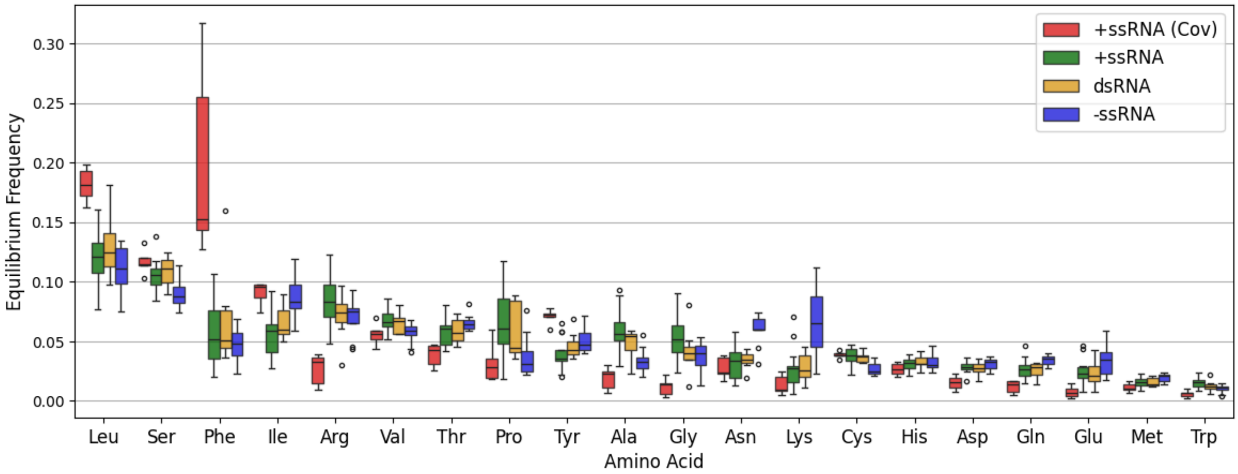


**Figure 13. Amino acid mutational equilibrium frequencies distributions for 4 groups of viruses.**

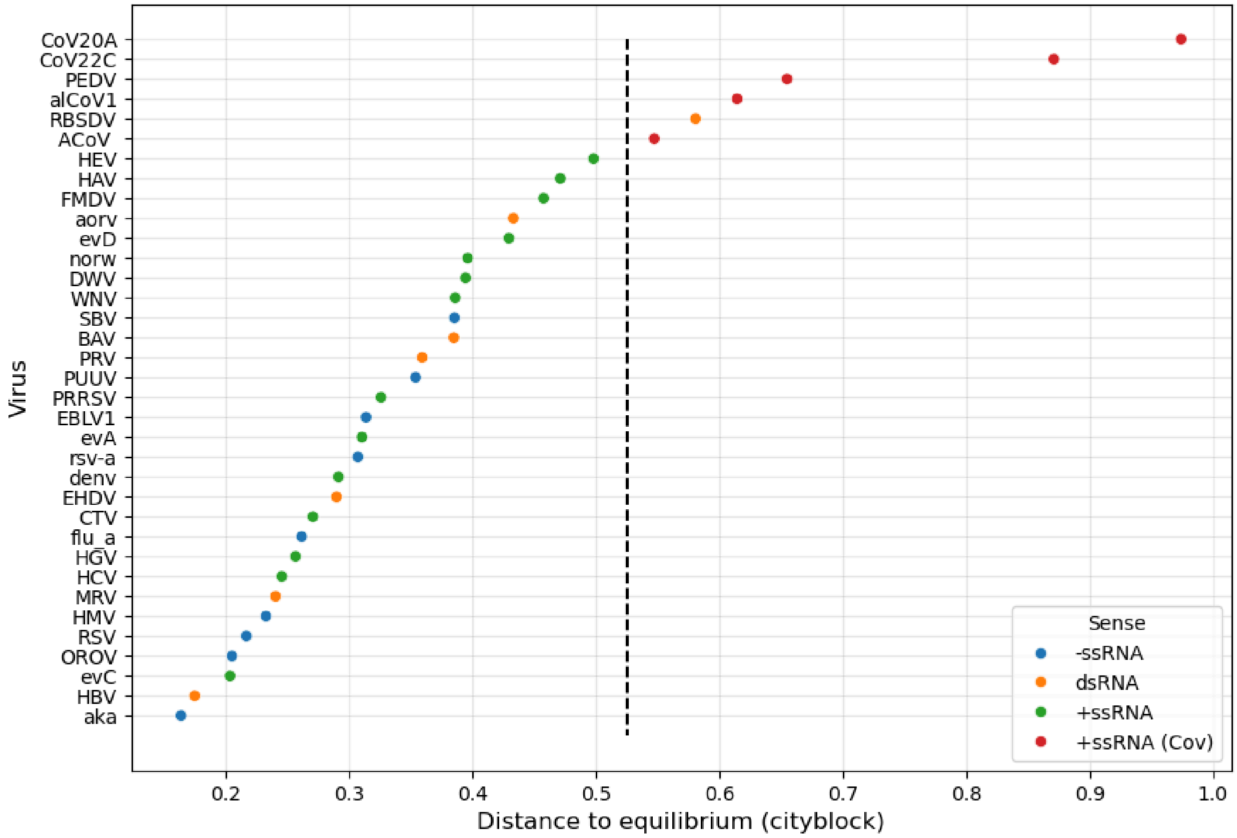


**Figure 14. Distance to equilibrium amino acid composition for viruses in our dataset.** The perpendicular marker denotes the boundary utilized for viral classification.

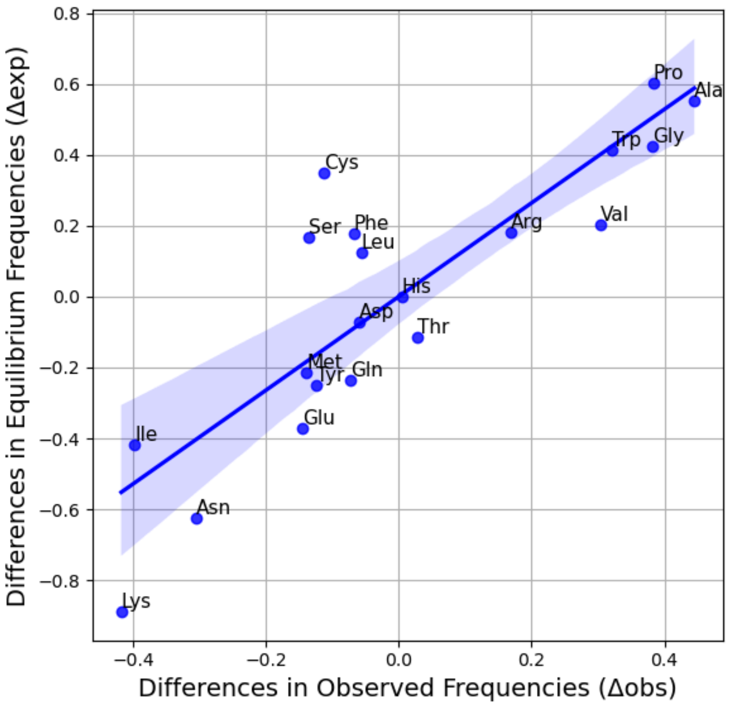


**Figure 15. Comparison of observed versus expected differences in amino acid composition between +ssRNA and –ssRNA viruses excluding distant from equilibrium viruses.**

Pearson r = 0.856, p = 1.5e-06, n = 20. Prior to correlation and subtraction, CLR-transformation was performed on the amino acid composition vectors. The average group-specific mutational spectra were derived from a cohort of 9 –ssRNA and 14 +ssRNA viral species, omitting representatives from the Coronaviridae family (refer to Fig. S13).

We observed that the alignment between expected and empirical compositional shifts reaches peak fidelity when viral genomes reside in proximity to their mutational equilibrium. Conversely, viral lineages situated at a distance from equilibrium—presumably due to recent shifts in their mutational spectrum—generate stochastic noise that degrades the observed correlation.

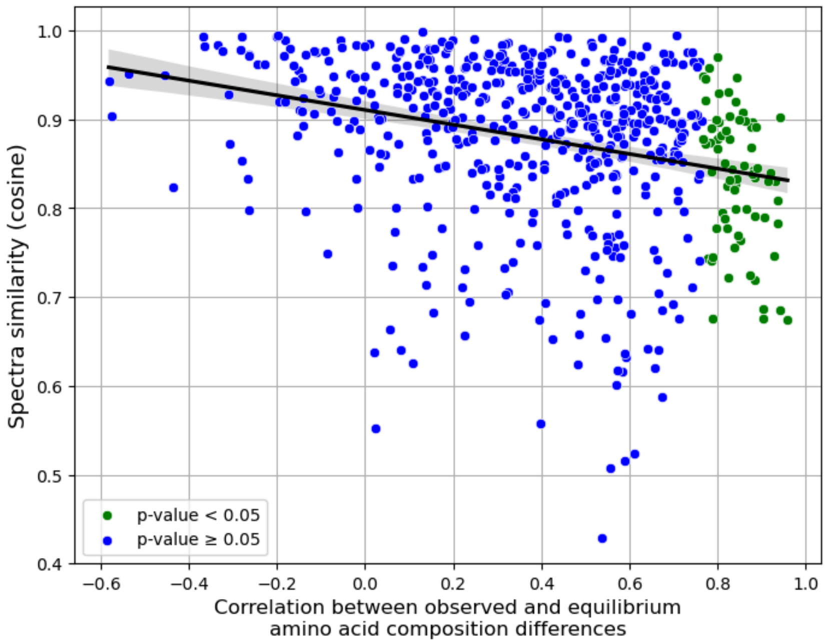


**Figure 16. Pairwise comparison of observed and expected differences in amino acid content among 34 viral species.**
CLR-transformation was applied to amino acid composition vectors before subtraction. Bonferroni correction was applied to the Pearson correlation p-values.

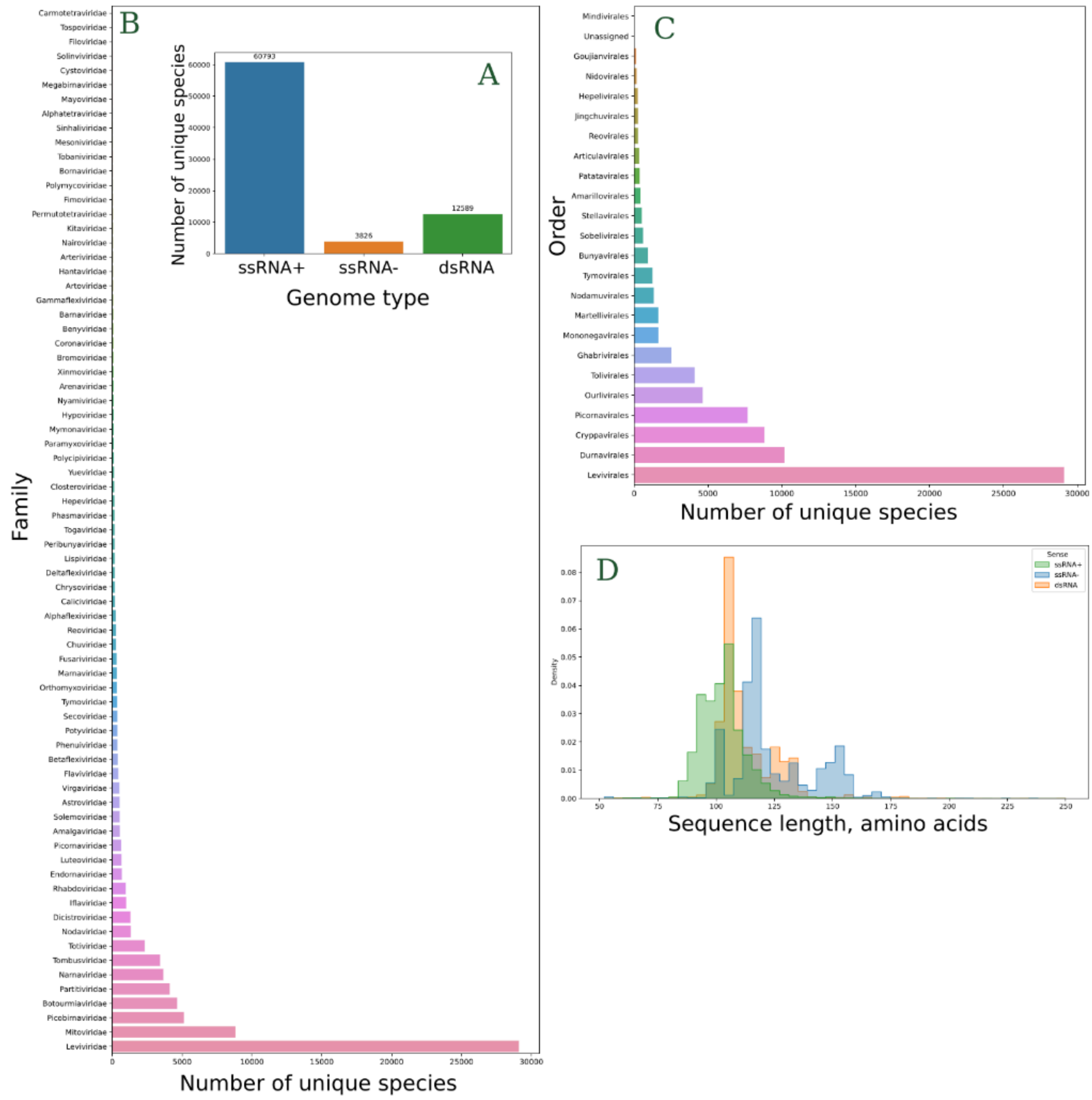


**Figure 17. Comprehensive overview of RNA virus species RdRp sequence dataset.**
(A) Species distribution categorized by genome sense; (B) representation of species counts across different families; (C) representation of species counts across different orders; (D) a summary of RdRp sequence length variations.

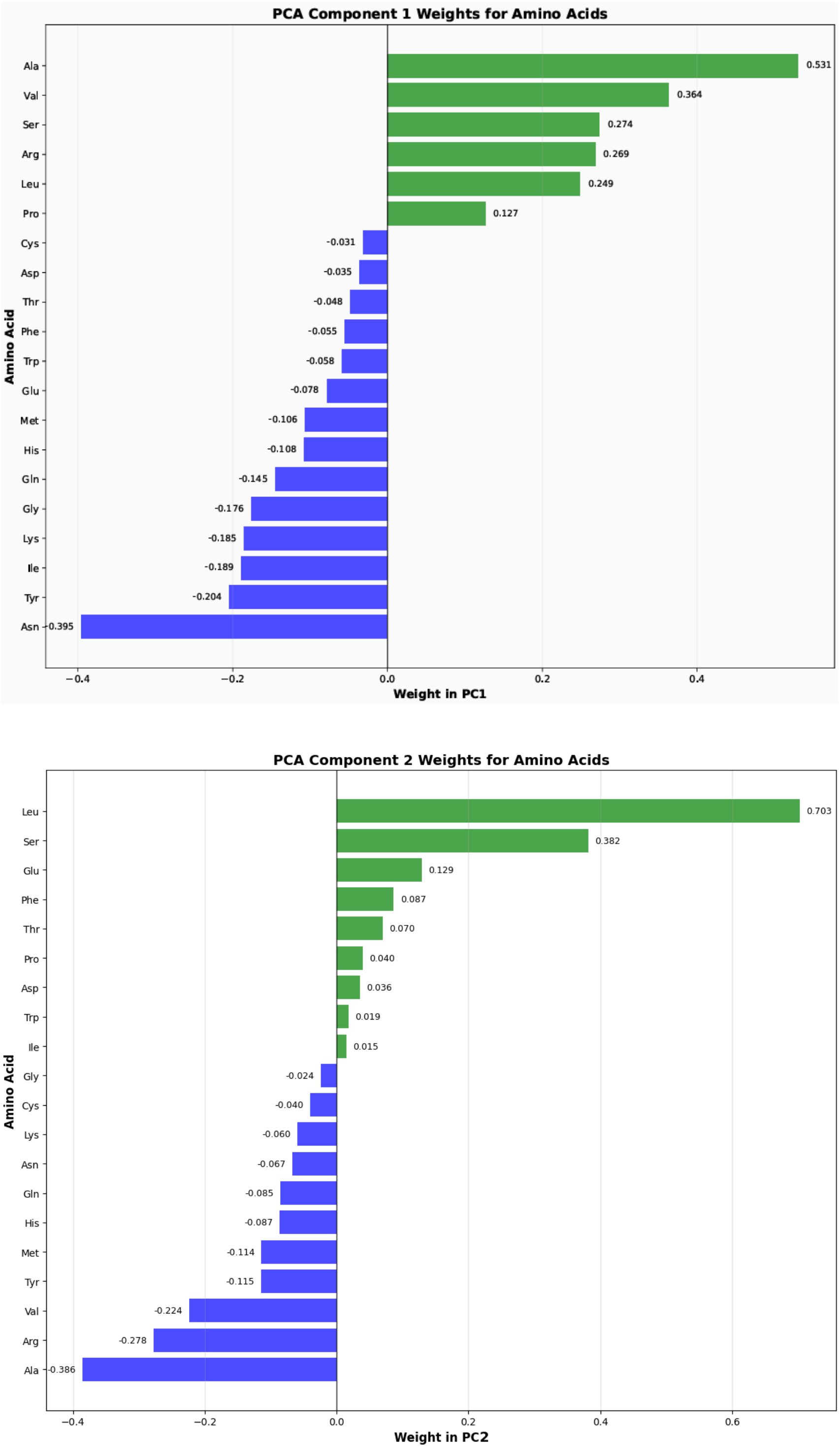


**Figure 18. The amino acids loadings in the first and second principal components of RdRp amino acid composition PCA-analysis.**
